# Global effects of focal brain tumours on functional complexity and network robustness

**DOI:** 10.1101/142844

**Authors:** Michael G Hart, Rafael Romero-Garcia, Stephen J Price, John Suckling

**Affiliations:** Brain Mapping Unit, Department of Psychiatry, Sir William Hardy Building, Downing Street, Cambridge, CB2 3EB; Academic Division of Neurosurgery, Department of Clinical Neurosciences, Box 167, Cambridge Biomedical Campus, Cambridge, CB2 0QQ; Brain Mapping Unit, Department of Psychiatry, Herchel Smith Building for Brain and Mind Sciences, Robinson Way, Cambridge, CB2 0SZ

**Keywords:** fractal analysis, Hurst exponent, resting state fMRI, glioblastoma

## Abstract

**Background:** Neurosurgical management of brain tumours has entered a new paradigm of supra-marginal resections that demands thorough understanding of their peri-tumoural functional effects. Historically the effects of tumours have been considered as local, and long-range effects have not been considered. This work tests the hypothesis that focal tumours affect the brain at a global level producing long-range gradients in cortical function.

**Methods:** Resting state functional (f)MRI data were acquired from 11 participants with glioblastoma and split into discovery and validation datasets. Fractal complexity was computed with a wavelet-based estimator of the Hurst exponent. Distance related effects of the tumours were tested with a tumour mask dilation technique and parcellation of the underlying Hurst maps. Functional connectivity networks were constructed and validated with different parcellation and statistical dependency methods prior to graph theory analysis.

**Results:** Fractal complexity, measured through the Hurst exponent and tumour mask dilation technique, demonstrates a penumbra of suppression in the peri-tumoural region. At a global level, as distance from the tumour increases, this initial suppression is balanced by a subsequent over-activity before finally normalizing. These effects were best fit by a quadratic model, and were consistent across different network construction pipelines. The Hurst exponent was significantly correlated with multiple graph theory measures of centrality including network robustness, but graph theory measures did not demonstrate distance dependent effects.

**Conclusions:** This work provides evidence to support the theory that focal brain tumours produce long-range and non-linear gradients in function. Consequently the effects of focal lesions, and the resultant clinical effects, need to be interpreted in terms of the global changes on whole brain functional complexity and network architecture rather than purely in terms of functional localisation. Determining whether the peri-tumoural changes represent adaption or potential plasticity, for example, may facilitate extended resection of the tumour without functional cost.

## INTRODUCTION

A new philosophy of supra-marginal resection has entered the vocabulary of neuro-oncology whereby surgical excision is extended beyond the confines of a lesion as it is visible on standard structural imaging. This paradigm is based on identification of tumour infiltration beyond the contrast enhancing margin (Miwa *et al*., 2004; Price and Gillard, 2011; Price *et al*., 2007; Witwer *et al*., 2002), fluorescence-guided imaging that can identify potentially infiltrated brain at surgery (Floeth *et al*., 2011; Stummer *et al*., 2006), clinical studies correlating extended resections with improved survival (Aldave *et al*., 2013; Ius *et al*., 2011; Li *et al*., 2016; Yan *et al*., 2017), and potentially improved effectiveness of adjuvant therapies (Stummer *et al*., 2012). However, extending surgery outside the lesion margins comes with the risk of intruding on eloquent brain and consequently the potential of neurological impairment. Therefore, understanding the function of the peri-tumoural brain is of paramount importance.

This requirement to accurately map brain function led to many notable contributions from neurosurgery in understanding functional neuroanatomy (Greenblatt *et al*., 1997; Penfield and Rasmussen, 1950). However, the traditional approach to understanding the effects of focal tumours *per se* on brain function is somewhat limited. Firstly, the effects of brain lesions on the surrounding tissue are believed to be only local (i.e. through structural disruption), and potential long-range effects are overlooked. Furthermore, function itself is viewed as being uniform and consequently represented as a binary outcome (i.e. brain either is, or is not eloquent); considering function as a continuous variable has not been investigated. Understanding putative long-range effects of brain tumours and non-binary gradients in function may allow more accurate prediction of the effects of tumour removal and insight into mechanisms of plasticity-related recovery.

Fractal analysis of resting state functional (f)MRI data offers the potential to understanding putative long-range gradients in brain function. Fractals are signals that display scale-invariance; that is, they have similar features regardless of the scale at which they are viewed. Fractal signals are pervasive in nature (examples include coastlines, snowflakes, and ocean waves) and in the brain (neuronal membrane potentials, neural field potentials as well as EEG, MEG, and blood oxygenation level dependent (BOLD) contrast endogenous signals) (He, 2014). From a fractal perspective, healthy homeostasis emphasises the complexity of the underlying biological processes rather than regular steady-state behaviour (Goldberger *et al*., 2002; Ivanov *et al*., 1999). In the brain this complexity is believed to enable adaptation to stimuli or other challenges, while simpler and less complex dynamics are indicative of less advantageous function or disease (Ivanov *et al*., 2001; Maxim *et al*., 2005; Suckling *et al*., 2008; Wink *et al*., 2008).

Connectomics is a novel multi-disciplinary paradigm that naturally views the brain as a complex network of individual components interacting through continuous communication, and offers a window into the effects of focal tumours at the global level (Hart, Ypma, *et al*., 2016). Furthermore, graph theory analysis of brain networks unlocks a new terminology with which to describe brain functional neuroanatomy and for predictive modelling of, for example, the effects of tumour removal and how this is related to network robustness (Hart, Price, *et al*., 2016). Combining fractal and network analysis methods provides a complimentary approach to identifying putative wide-ranging effects of focal brain tumours on signal complexity and network robustness.

In this work, the concept that tumours only lead to local effects on brain function is challenged and a new theory that considers the global effects of a tumour is proposed that can augment our current brain mapping techniques. BOLD sensitive, resting-state fMRI was used to map brain function with data-driven analysis of fractal complexity and functional connectivity at the whole brain level. It was hypothesized that tumours would produce long-range effects outside of their margin as depicted on structural MRI, and that there would be gradients of BOLD signal and functional connectivity extending away from the tumour. Evidence is presented to support the theory that focal brain tumours produce long-range and non-linear gradients in function.

## MATERIALS AND METHODS

### Experimental design

The study was approved by the Local Regional Ethics Committee (protocol number10/H0308/23) and all participants provided written informed consent. Inclusion criteria were the MRI appearance of a tumour consistent with a glioblastoma. In fact, all participants had a confirmed glioblastoma at local histological review according to WHO criteria (Louis *et al*., 2016).

We selected a cohort of five participants to form a discovery dataset for hypothesis generation. Subsequently, a further six participants were included, forming the validation dataset. The discovery dataset was designed to be homogenous and included moderately sized tumours based on the right parietal lobe, whereas the validation dataset was designed to be more heterogeneous with a variety of tumour sizes and locations. Splitting the group into separate discovery and validation cohorts was performed to test if findings were replicable across a variety of tumour presentations. Demographic information is summarized in table 1.

**Table 1:** Participant demographics. Extent of resection is based on RANO criteria [Wen et al, 2010]. Molecular markers include MGMT promoter methylcation, IDH-1 mutation, and 1p19q status. MIB-1 = mindbomb antibody. *censored data

| Age | Sex | Seizures | Pre-op exam | Post-op exam | Pathology | Surgery | Tumour location | Molecular markers | Survival (months) |
| --- | --- | --- | --- | --- | --- | --- | --- | --- | --- |
| 64 | F | Pre-op | Left pronator drift | Homonymous hemianopia | Glioblastoma | No residual disease | Right superior parietal lobule | Negative / 85% | 23 |
| 73 | F | No | Intact | Homonymous hemianopia | Glioblastoma | No residual disease | Right inferior parietal lobule to occipital pole | Negative / MIB 32% | 13 |
| 79 | M | No | Hemianopia | No change | Glioblastoma | No residual disease | Right inferior occipital lobe | Negative / MIB 28% | 17 |
| 76 | M | Pre-op | Left hemiparesis | Contralateral motor paresis deterioration | Glioblastoma | Biopsy | Right superior para-central lobule | Negative / MIB 18% | Unavailable |
| 36 | M | Post-op | Left hemiparesis | No change | Anaplastic astrocytoma | No residual disease | Right post-central gyrus and supramarginal gyrus | Negative / MIB 30% |  |
| 62 | M | No | Hemianopia | No change | Glioblastoma | Biopsy | Right thalamic extending to temporo-parietal-occipital region | Negative / MIB 12.5% | 14* |
| 52 | M | No | Left hemiparesis | No change | Glioblastoma | Complete resection | Right supramarginal gyrus | Negative / MIB 25% | 15* |
| 57 | M | No | Memory & language | No change | Glioblastoma | Complete resection | Left superior frontal gyrus | Negative / MIB 26.1% | 16* |
| 36 | M | No | Hemianopia | No change | Glioblastoma | Complete resection | Right parieto-occipital | Negative / MIB 8.3% | 18* |
| 44 | M | Pre-op | Nil | No change | Glioblastoma | Biopsy | Left caudate | IDH-1 & MGMT positive / MIB 38% | 12* |
| 50 | M | No | Hemianopia | No change | Glioblastoma | Complete resection | Right supramarginal gyrus / occipital lobe | Negative / MIB 45% | 8 |

### Imaging parameters

MRI data were acquired using a Siemens Trio 3T scanner and 16-channel receive-only head coil (Siemens Medical Solutions). A multi-echo echo planar imaging sequence was performed for 10 minutes and 51 seconds at a repetition time, TR = 2.42s resulting in 269 three-dimensional volumes covering the cerebral cortices and cerebellum. Acquisition parameters for resting fMRI were: flip angle = 90 degrees; matrix size = 64x64; in-plane resolution = 3.75mm; echo times (TE) = 13.00ms, 30.55ms, 48.10ms; slice thickness = 3.8mm. Anatomical images were acquired using a T1-weighted magnetization prepared rapid gradient echo (MPRAGE) sequence (FOV = 256mm x 240mm x 176mm; matrix = 256x240x176; voxel size = 1mm isotropic; TR = 2300ms; TE = 2.98ms; flip angle = 9 degrees).

### Structural MRI pre-processing

A summary of the analysis pipeline is presented in figure 1. Advanced Normalization Tools (ANTs) cortical thickness pipelines (Avants *et al*., 2009; Das *et al*., 2009; Tustison *et al*., 2013) were used to perform automated volume based cortical thickness estimation, the workflow including: initial N4 bias correction on input anatomical MRI; brain extraction using a hybrid segmentation/template-based strategy; alternating between prior-based segmentation and white matter posterior probability weighted bias correction; DiReCT-based cortical thickness estimation in native space. A hand drawn tumour mask to encompass the contrast-enhancing margin was created by the neurosurgeon and used as an additional prior for segmentation. Finally, images were non-linearly mapped to MNI space using symmetrical diffeomorphic registration with masking of the tumour from cost-function calculation(Andersen *et al*., 2010).

**Figure 1:**
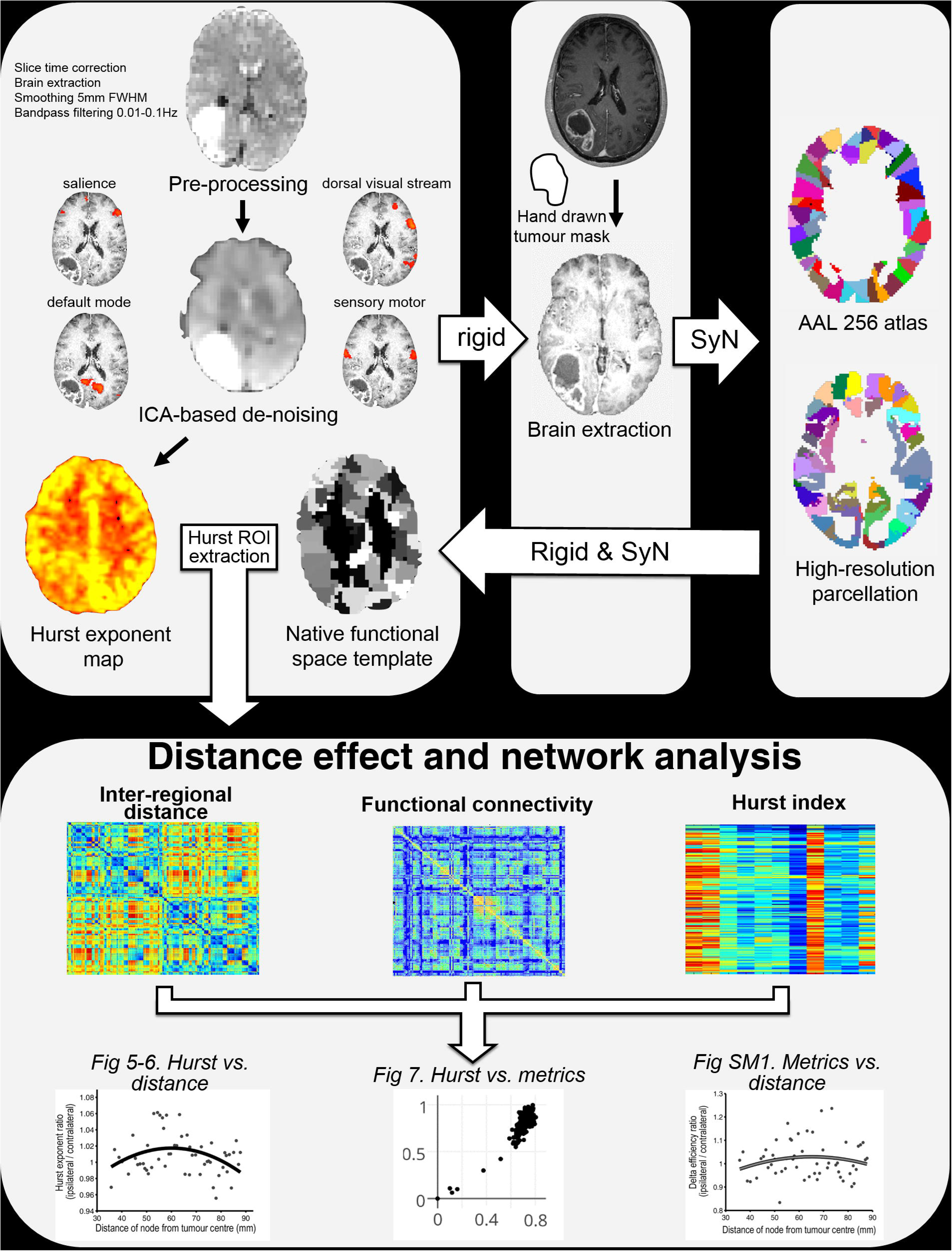
structural and functional image analysis pipeline. Functional images underwent multi-echo ICA-based de-noising and selected pre-statistical processing in functional space. Anatomical images underwent brain extraction, bias field correction, and tissue segmentation (including hand drawn masks of the tumour contrast-enhancing volume). Standard space parcellation templates were warped to the structural image using symmetrical diffeomorphic registration and a binary tumour mask to exclude the region from cost function weighting. An additional rigid 6 degree of freedom transform was performed from structural to functional space, then these transforms were combined to produce a representation of the parcellation template in functional space, obviating the need for interpolation of the functional image. Fractal analysis was performed in structural space to allow optimal delineation of the tumour mask. SyN = symmetrical diffeomorphic normalisation. FWHM = fixed width at half maximum.

### Resting state fMRI pre-processing

Data pre-processing was performed using AFNI (Cox, 1996) (http://afni.nimh.nih.gov/afni/). The first 15 seconds of image data were discarded to allow for magnetization to reach steady state. Subsequent steps included: slice time correction; rigid-body motion correction; de-spiking; and functional imaging brain extraction. For de-noising, multi-echo independent component analysis (ME-ICA) was used (Kundu *et al*., 2012; 2013). Linear registration matrices (6 degrees of freedom) between functional and structural acquisition space were calculated with ANTs.

### Fractal analysis

Fractal analysis was performed with a wavelet based estimator of the Hurst exponent (figure 2) (Bullmore *et al*., 2003; 2004; 2009) applied to processed resting state fMRI data registered to the corresponding structural image. In brief, time series at each voxel underwent wavelet decomposition. A linear model was regressed on to the logarithm of the variance of wavelet coefficients as a function of scale. The estimated slope of this model is proportional to the Hurst exponent, H, and in turn the fractal dimension. Hurst exponents were obtained for every intracerebral voxel, and then thresholded at H<0.5 (corresponding to white noise) and values were extracted for each parcel of the parcellation template (see below).

**Figure 2:**
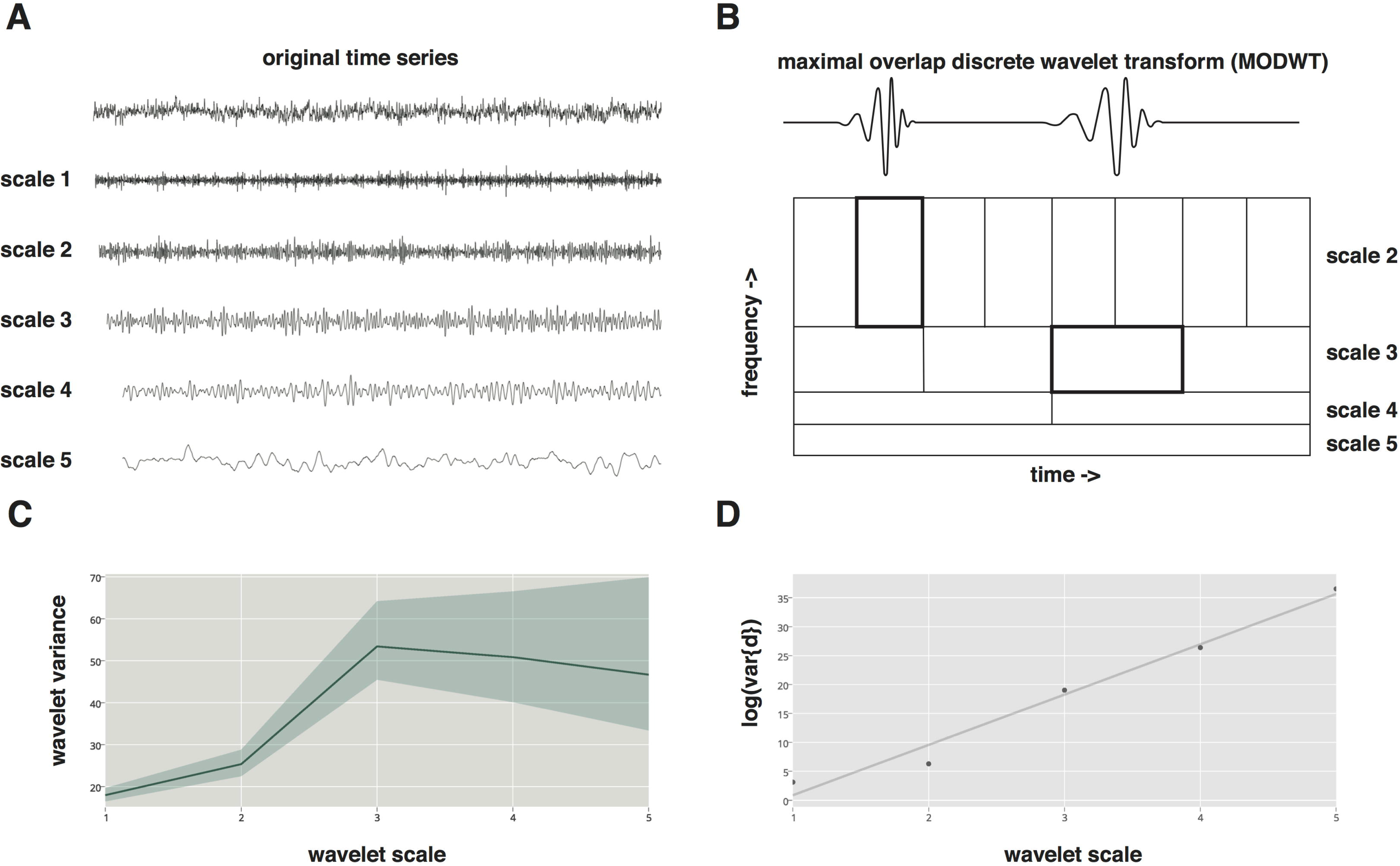
fractal analysis and hurst exponent calculation. A. The time series of BOLD contrast for a selected brain region (in this case the right superior frontal gyrus) is displayed on top: underneath are the same time series at wavelet scales one to five. B. The maximal overlap discrete wavelet transform (MODWT) scales the wavelet in frequency and amplitude to decompose the original time series at multiple scales (note the planned decomposition is also a natural fractal). Here we illustrate Daubechies 8 wavelet (note it integrates to 0 and has an irregular shape) at two scales, above. C. Wavelet variance of the decomposed time series scales plotted against scale. As wavelet scale increases (and therefore frequency decreases) the variance increases. D. Self-similarity of variance is exponentially related to the scale and therefore forms a straight line on a plot of wavelet scale versus the log of the variances of the wavelet (detail) coefficients at the corresponding scale, log(var{d}). The exponent of the straight line is related to the Hurst exponent 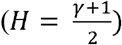 and thereafter the fractal dimension (*D* = 2 – *H*).

### Peri-tumoural brain analysis

The hand drawn binary tumour masks were dilated to 30mm beyond their original margins in 2mm increments, followed by subtraction of the preceding masks. This created a series of 2mm annuli expanding from the tumour margin. These annuli were then masked, only being included in further analyses where they overlapped the cortical grey matter estimated from the structural MRI (figure 3). For each mask, for each participant, average H estimated from fMRI was extracted for each annular mask. Masks were subsequently reflected across the interhemispheric fissure to compute the corresponding values from the contralateral, healthy hemisphere. Values were then expressed as the ratio of H in homologous regions.

**Figure 3:**
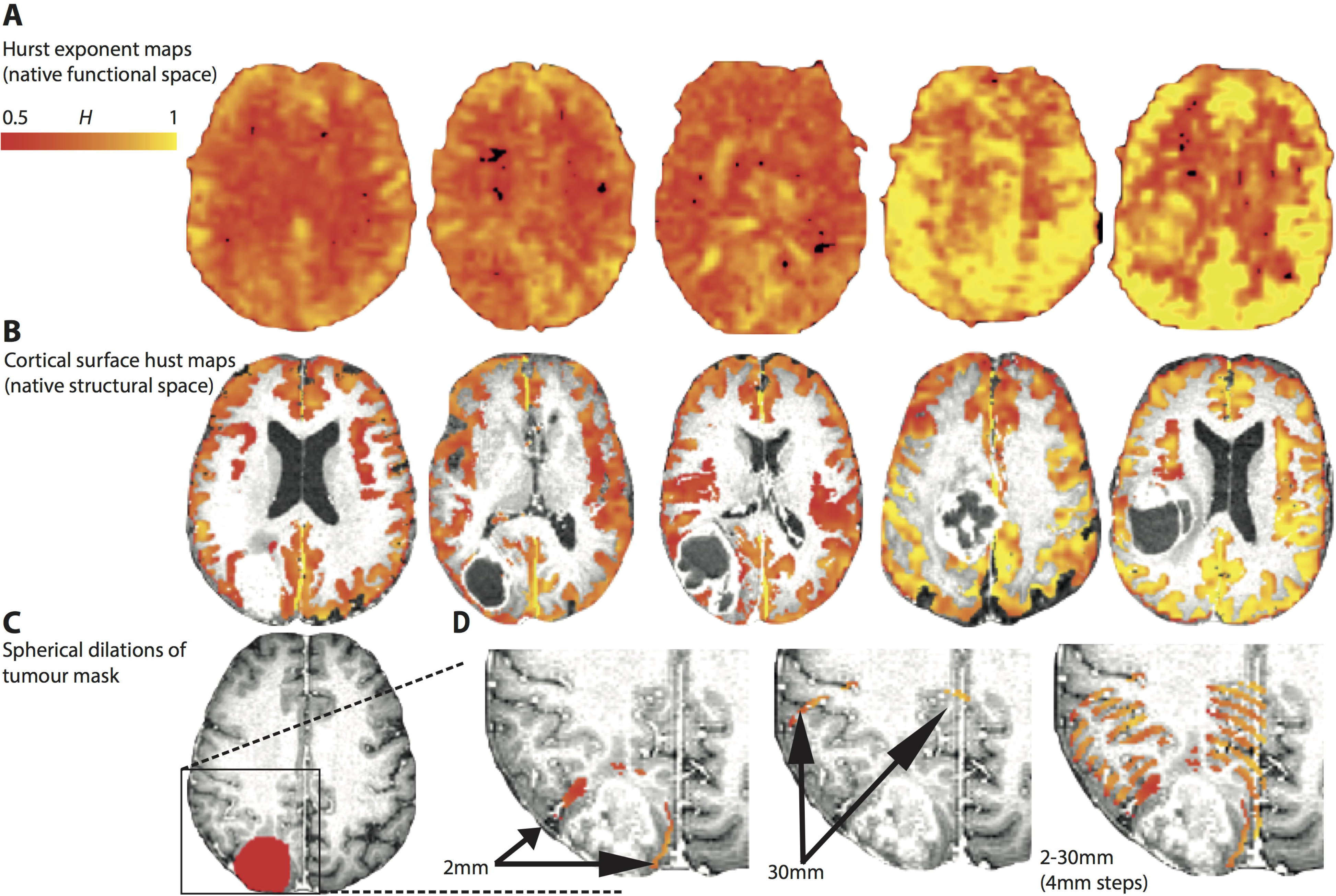
sequential mask dilations. A: Hurst exponent maps (3D volumes) for each patient in the discovery cohort. B: Brain extracted and intensity normalized structural images corresponding to each individual. Overlaid are the cortical thickness maps (generated in ANTS) that are used to mask the raw Hurst exponent maps (in structural space) i.e. the Hurst exponent is extracted specifically for the grey matter. C: A detail from a single patient with an example binary tumour mask drawn in structural space. D: The tumour mask subsequently underwent sequential 2mm dilations with subtraction of the previous tumour mask to produce a 2mm thick tumour rim. This dilating tumour rim is used to extract H values within the cortex. E: The image on the right shows the results of dilations at 2, 6, 10, 14, 18, 22, 26, and 30 mm.

### Connectome analysis

A summary of the connectome analysis pipeline is presented in figure 1. Parcellation was initially performed with an anatomical template of 251 equal sized parcels (Romero-Garcia *et al*., 2012) then subsequently all analyses were repeated on a randomized template of 256 equal sized parcels (Zalesky *et al*., 2010) to test for parcellation independence of network features. Parcels overlapping the tumour mask were removed from subsequent processing. Statistical dependencies between mean processed time series of each parcel were computed using wavelet, Pearson, and partial correlations to test for independence of network features from the underlying method of connectome constructions. Negative correlations were excluded but thresholding was not applied (i.e. matrices were ‘fully connected’).

A graph is composed of nodes (brain regions) and links, which are associated with weights that denote the strength of the connection between nodes. To reduce multiple comparisons and repeated analysis of non-independent network measures, specific graph theoretical measures were specified *pre hoc*, chosen to reflect fundamental centrality features of the underlying network: node strength (the sum of weights to a node); global efficiency (the average inverse shortest path between nodes); betweenness centrality (identifying nodes that participate in the shortest paths (Freeman, 1978)); within-module degree z-score (identifying nodes that are highly integrative within individual modules, but not necessarily between modules (Guimera and Amaral, 2005)); participation coefficient (identifying nodes that play a key role linking multiple distinct modules, the converse of the within-module degree z-score (Guimera and Amaral, 2005)); eigenvector centrality (the average centrality of a node’s neighbours (Bonacich, 1972)).

A network measure particularly pertinent to neurosurgery is node robustness. This is tested by synthetic lesioning, whereby a node (and all of its connections) is removed from the network and the percentage change (‘delta’) of a selected global network measure is computed. Typically this is performed with global efficiency in which case the resultant measure is also known as delta efficiency. Each node’s delta efficiency values were expressed as a ratio over the value in the homologous node in the contralateral hemisphere. Euclidean distance was defined from the centre of each node to the centre of the tumour. Network measures were computed using the brain connectivity toolbox (Rubinov and Sporns, 2010) and transformed to Z-scores.

### Statistical analysis

Between-tissue differences in H were tested with ANOVA, with the threshold for significance, p <0.05. Testing relationships with distance for the dilating tumour mask and H parcellation analyses was performed with linear, quadratic, and cubic models. Goodness of fit was determined with the Akaike Information Criteria (Akaite, 1974). All correlations between network measures underwent Bonferroni correction for multiple comparisons.

## RESULTS

### 1. Tissue Hurst values

Whole-brain Hurst exponent maps, and H masked by cortical segmentation estimated with ANTs from the structural MRI of the discovery cohort, are presented in figure 3. Brain extraction, registration and segmentation with ANTs resulted in high quality images using default parameters without noticeable artefacts due to the presence of the tumour. Differences in the means of the Hurst exponent between neocortical grey matter and white matter for the discovery, validation, and complete cohorts were not significant (ANOVA, F = 1.28, p = 0.31, figure 4), although the observed higher values in grey, relative to white matter are consistent with the previous findings (Wink *et al*., 2008).

**Figure 4:**
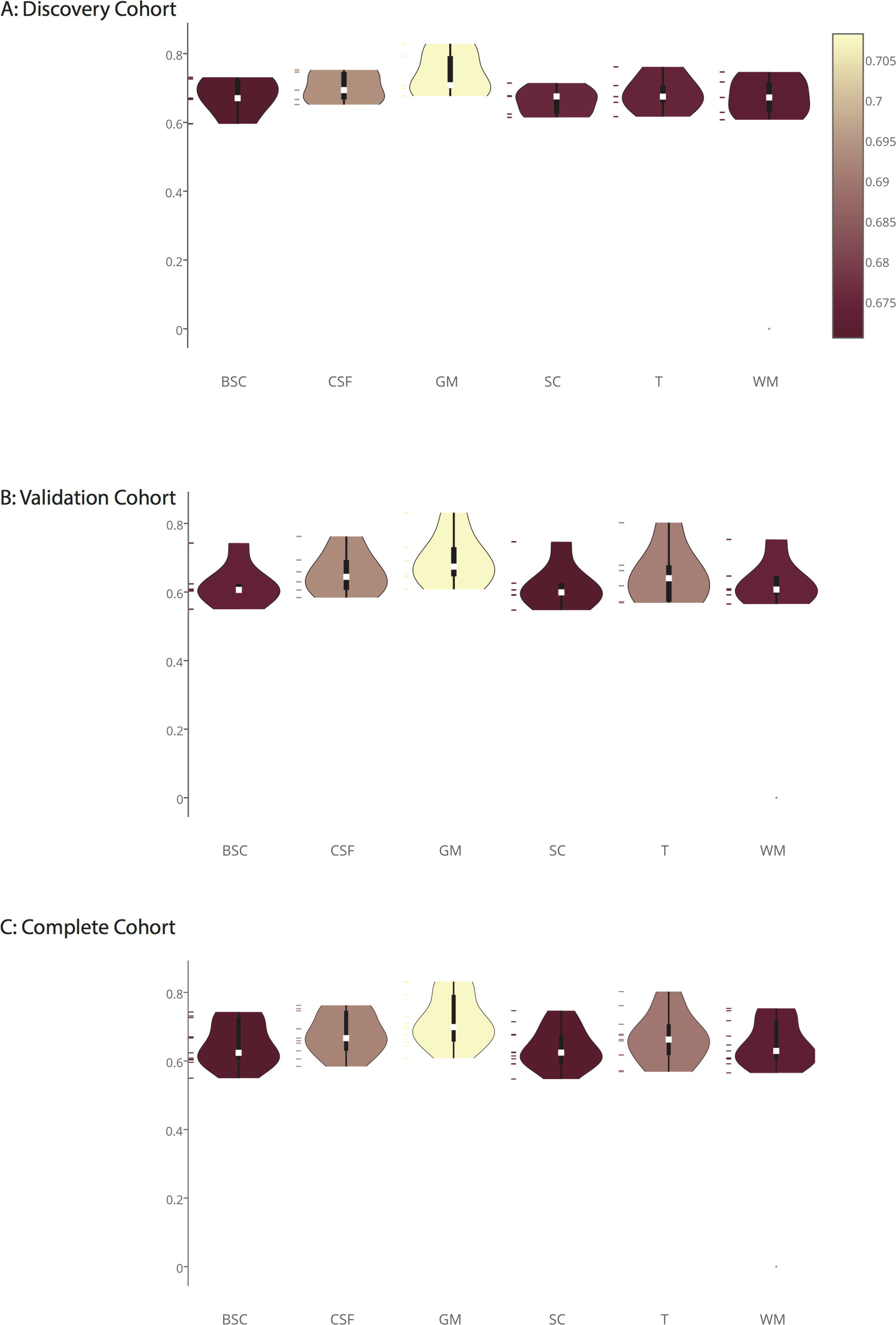
Hurst tissue segmentations. Box plots of extracted H group mean values for tissue segmentations including grey matter, white matter, subcortical structures, contrast enhancing tumour volume, and infratentorial structures (brain stem and cerebellum). Median H is denoted by the white square and colour coded, boxes represent interquartile ranges, and whiskers represent 1.5 times the interquartile range. BSC = brainstem and cerebellum, CSF = cerebrospinal fluid, GM = grey matter, SC = subcortical nuclei, T = tumour, WM = white matter.

### 2. Peri-tumoural Hurst effects with distance

Using the tumour mask sequential dilation technique revealed a quadratic increase in H with distance from the tumour (R^2^ 0.6−0.77, p<0.01) in the discovery cohort. Both validation and complete study cohorts also demonstrated this quadratic increase in H with distance from the tumour demonstrating the replicability of the finding (figure 5). A penumbra of reduced complexity relative to the contralateral hemisphere extended for 10-15mm from the tumour border while beyond 15mm complexity continues to increase beyond the values seen contralaterally, albeit non-monotonically, suggesting a complex relationship to global changes.

**Figure 5:**
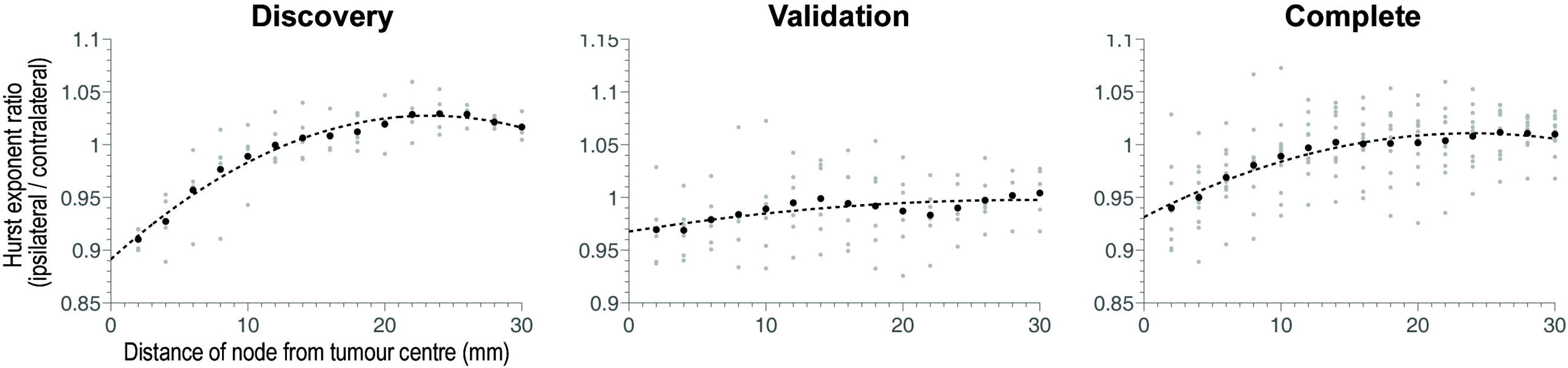
Hurst exponent versus distance from tumour centre in dilating tumour masks. The mean H value is plotted as a function of distance from the tumour border in 2mm increments up to a maximum of 30mm. The cohort is split into discovery, validation, and complete cohorts from left to right, all of which demonstrate significant quadratic fits (table 2).

### 3. Whole brain parcellation of Hurst versus distance

To expand on the changes in H related to the distance from the tumour border, the H maps were parcellated using the same templates as for the connectome analysis, with the value of each parcel reflecting the mean H. This allowed greater coverage and a whole brain analysis to complement the limited range of the dilating spherical mask, but with coarser resolution than the dilating tumour mask approach. Results were consistent with those from the tumour mask sequential dilation technique in that closest to the tumour values were reduced compared with the contralateral hemisphere, then increased with distance to above contralateral levels, before finally reducing again back to contralateral values (figure 6). This trend was best fit with a quadratic model (R^2^ 0.07−0.12, AIC −9.77 to −12.2; table 2) in the discovery cohort. The validation cohort also demonstrated this quadratic variation in H with distance from the tumour, again demonstrating replicability as with the sequential mask dilation technique. Furthermore, results were independent of the parcellation technique, with both anatomical and randomised parcellation templates demonstrating the same quadratic variation in H with distance from the tumour in all three cohorts (discovery, validation, and complete).

**Figure 6:**
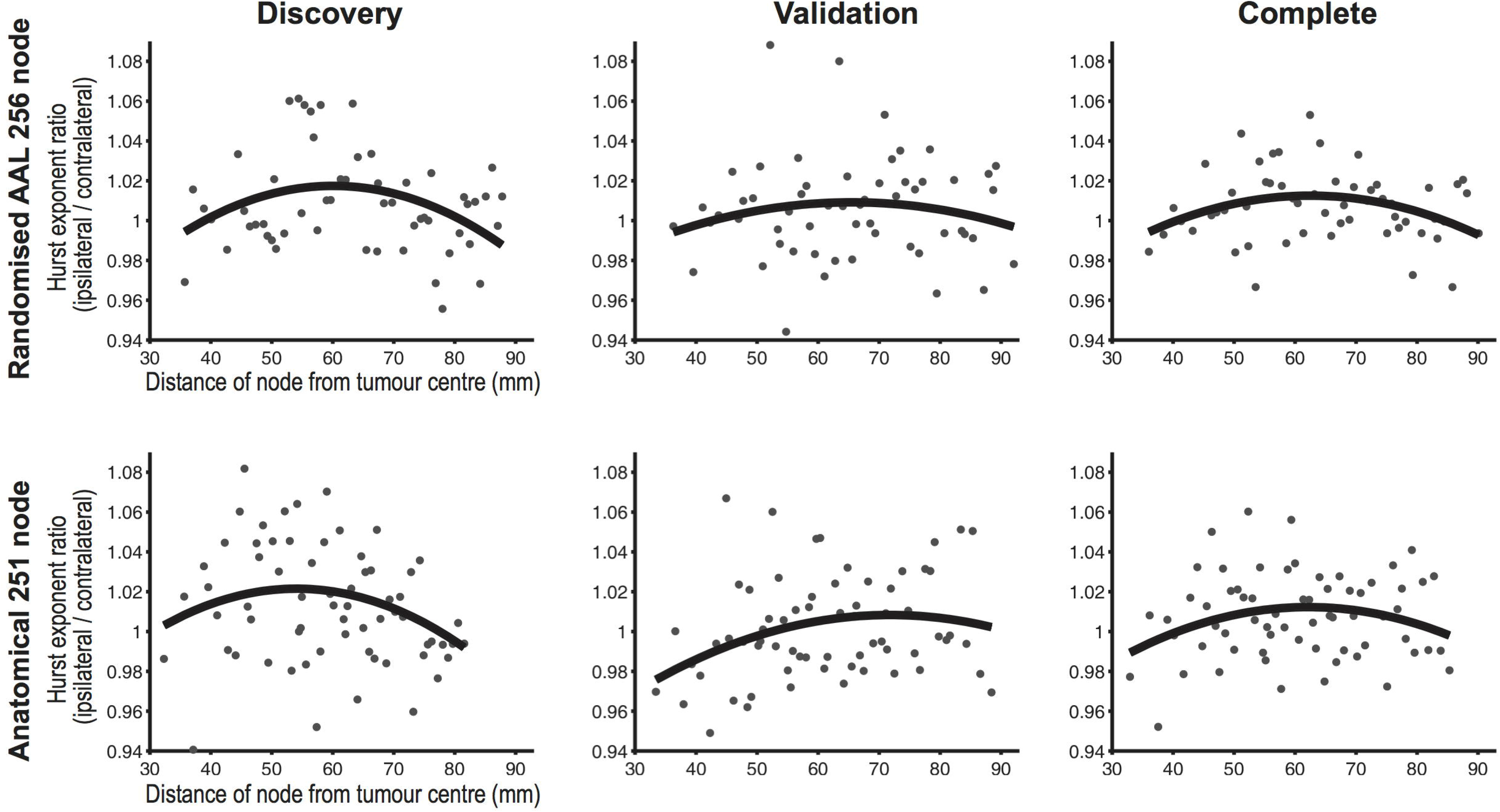
Hurst exponent versus distance in parcellation nodes. The ratio of H between homologous parcels in each hemisphere is plotted against distance from the tumour centre. Both parcellation templates are represented (top row = randomised, lower row = anatomical). Participants are split into discovery, validation, and complete cohorts from left to right. The quadratic model demonstrated the best fit for all plots and is highlighted.

**Table 2:** Hurst parcellation distance effects. The best fit for discovery, validation, and complete cohorts for both parcellations was the quadratic model as determined by R^2^ and AIC, and highlighted in bold. AIC = Akaike information criteria

|  | Anatomical parcellation |  |  |  |  |  | Randomised parcellation |  |  |  |  |  |
| --- | --- | --- | --- | --- | --- | --- | --- | --- | --- | --- | --- | --- |
|  | Discovery |  | Validation |  | Complete |  | Discovery |  | Validation |  | Complete |  |
| *Fitting* | $R^2$ | *AIC* | $R^2$ | *AIC* | $R^2$ | *AIC* | $R^2$ | *AIC* | $R^2$ | *AIC* | $R^2$ | *AIC* |
| Linear | 0.02 | -9.70 | 0.056 | -10.48 | 0.00 | -11.15 | 0.02 | -10.80 | 0.00 | -10.55 | 0.00 | -12.04 |
| **Quadratic** | **0.07** | **-9.77** | **0.085** | **-10.51** | **0.06** | **-11.23** | **0.12** | **-10.99** | **0.02** | **-10.56** | **0.10** | **-12.20** |
| Cubic | 0.06 | -9.72 | 0.084 | -10.48 | 0.05 | -11.19 | 0.11 | -10.92 | 0.02 | -10.52 | 0.09 | -12.15 |

### 4. Whole brain network measures distance effects

To further expand upon the distance related changes in H, we investigated whether functional connectome graph theory measures also demonstrated complimentary distance related effects, specifically that of network robustness. No significant distance related trends, linear or quadratic, were identified for any graph theory measure (**Figure_SM1**). Again, results were consistent across discovery and validation cohorts, anatomical and randomised parcellation templates, and with links weighted by either Wavelet or Pearson product-moment correlations.

### 5. Network measures relationship to Hurst exponent

Finally, given that node H, but not graph theory measures were related to distance from the tumour margin, the relationship between H of resting state BOLD time series and functional connectome graph theory measures was explored. H was significantly correlated (r = 0.33−0.87, p<10^−6^) with all selected network measures, although distributions were negatively skewed (Figure_SM2).

## DISCUSSION

### Summary

In this paper combined fractal and connectomic analyses were performed in a cohort of participants with brain tumours to determine whether the functional sequelae of focal lesions are apparent at the global rather than just the local level. Fractal scaling showed long-range gradients with distance from the tumour margin are best fit by a quadratic model that describes proximal suppression of complexity, followed by a subsequent increase, before final normalisation to values similar to those in the contralateral hemisphere. Furthermore, there was a correspondence between fractal properties of the BOLD signal and network measures, specifically those corresponding to centrality and network robustness. Together gives credence to the novel hypothesis that the brain responds to focal injury by reconfiguring its complexity globally, or at least far beyond the tumour margin. Previously, functional localisation has been based on the premise that focal injury is responsible for the resultant phenotype; this work suggests that the functional sequelae of focal brain tumours, and the resultant clinical effects, need to be interpreted in terms of spatially extended changes to whole-brain functional complexity and network architecture.

### Interpretation of findings

In the context of BOLD signal resting state functional MRI, a higher value of H represents increased complexity of the time series and better neuronal function (i.e. a more complex signal is able to carry more information and hence represents increased processing capacity). This has been corroborated in published studies commenting on the relationship between H and cognitive function, neurodegeneration, and pharmaceutical enhancement of performance (Maxim *et al*., 2005; Suckling *et al*., 2008; Weber *et al*., 2013; Wink *et al*., 2008). In this particular instance, the reduction in the Hurst exponent in the peri-tumoural region could represent a penumbra of regional cortical suppression, which may arise from direct tumour invasion or distant electro-chemical effects.

Correlation between centrality measures and network robustness implies the peri-tumoural suppression of H relates to brain tissue that has already had role reduction within the network. In terms of extended surgical resections, this suggests that removal of the surrounding peri-tumoural brain might be tolerated at the functional and network levels, and might not result in significant additional functional impairment. Conversely, if the tumour was mediating an electro-chemical suppressant affect, focused removal of only the lesion may ‘release’ this suppression and facilitate functional recovery. Key questions to be answered regarding the clinical effects of this phenomena are how it relates to function and impairment, and whether it recovers after removal of the contrast enhancing tumour mass.

### Interpretation of network measures

In this analysis, no distance related effects were found for any graph theory measure, but they were in turn significantly correlated with H. This discrepancy in distance related effects for H and network measures could of course be related to the lack of correspondence between them, or because the distance related effects themselves only partially explain the overall variance in the data. H is a local measure of BOLD signal complexity, and this appears to be altered in a way that is consistent across patients. However, these non-linear distant related changes to brain physiology may impact graph theoretical measures of centrality that characterize topological properties over spatially extended brain circuits in a more individual manner, diluting consistent distance related effects.

### Clinical applications at the individual level

Effects of focal cerebral lesions on cognition are determined by a number of factors (Anderson *et al*., 2009; Dennis, 2010; Opdebeeck *et al*., 2015; Stern, 2013), while detailed cognitive examination often reveals highly variable findings between patients despite broadly similar histological and demographic characteristics (Klein *et al*., 2012). Understanding the peri-tumoural brain at the individual level will be critical to determining the safety of resections, or indeed if it is possible to safely exceed the margins of a lesion in an effort to increase survival. Fractal analysis could be used to identify regions of suppressed cortical function that could be safely resected, or indeed areas that may improve upon resection of the main tumour volume and should be left untouched. However to perform reliable analyses at the individual level longitudinal imaging studies will be required to determine the dynamic network effects of the tumour and surgery over time.

### Further work and study limitations

This study is a proof-of-concept investigation into the effects of focal tumours on the complexity of BOLD signals and the functional connectome at the global level. While the overall number of participants were limited, it was felt the advantages of dividing the cohort into discovery and validation groups, namely increased methodological robustness and the ability to test replicability of findings, outweighed any marginal increase in statistical power from analysing participants in a single cohort. This resulted in relatively small cohorts, but with the benefit of each participant acting as their own control, this was not deemed to preclude analysis, particularly given the goal was to identify robust findings at the global level.

A rational follow-up to this work would be to include cognitive and longitudinal follow-up data to test the generated hypotheses; namely, that cognitive sequelae are dependent on the distance of resection from the tumour margin. Cognitive and longitudinal outcome data will aid inference on the compensatory or decompensatory nature of long-range functional gradients. Larger heterogenous cohorts will help determine whether the network effects are consistent or a feature of specific locations, and whether there is individual variation in the gradients of changes. Finally, these data also provoke discussion on the best principled parcellation strategies for focal tumours. Specifically, whether to include or exclude parcels either completely or partially overlapping the tumour, and possibly the surrounding area.

### Conclusion

Fractal and connectomic analyses suggest global rather than local effects of focal tumours on brain function. Furthermore, exploration of fractals and complex network measures revealed a strong relationship between signal complexity and graph theory measures, notably those reflecting network robustness. It is therefore conceivable that extent of resection, a key prognostic marker for survival, could be approached as an optimization problem balancing increased tumour removal with fractal scaling and network robustness. Further data could be used to test hypotheses that these distance-related effects reflect brain plasticity and potentially allow extended resection of the tumour without functional cost.

## ACKNOWLEDGEMENTS

Nil.

---

**Supplementary information figure 1:**
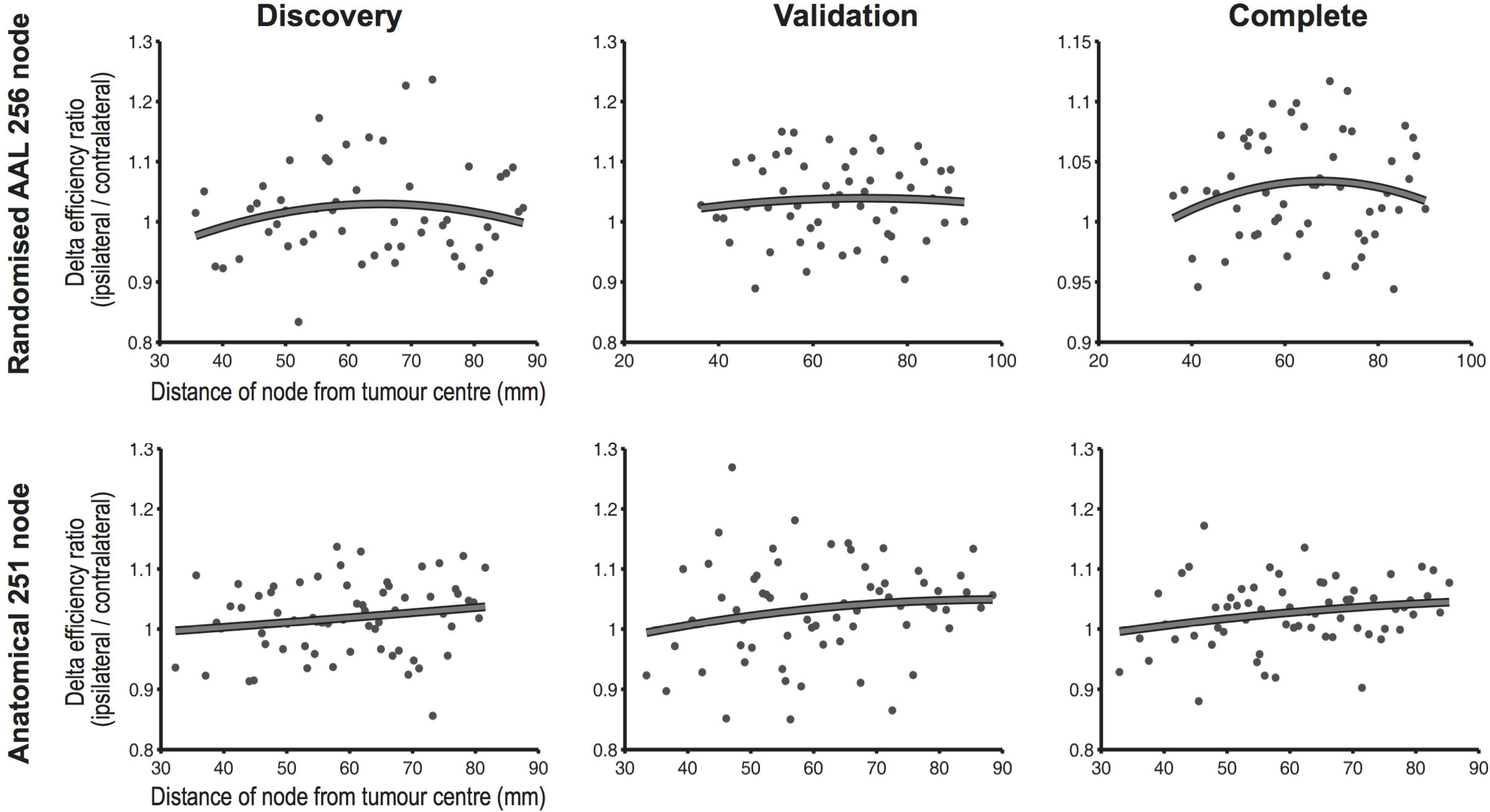
network efficiency versus distance. Top row images represent randomised parcellation template, lower row represents anatomical parcellation template, both for networks constructed with Pearson correlations. No consistent significant fits were demonstrated. Similar results are present with wavelet and partial correlation networks.

**Supplementary information figure 2:**
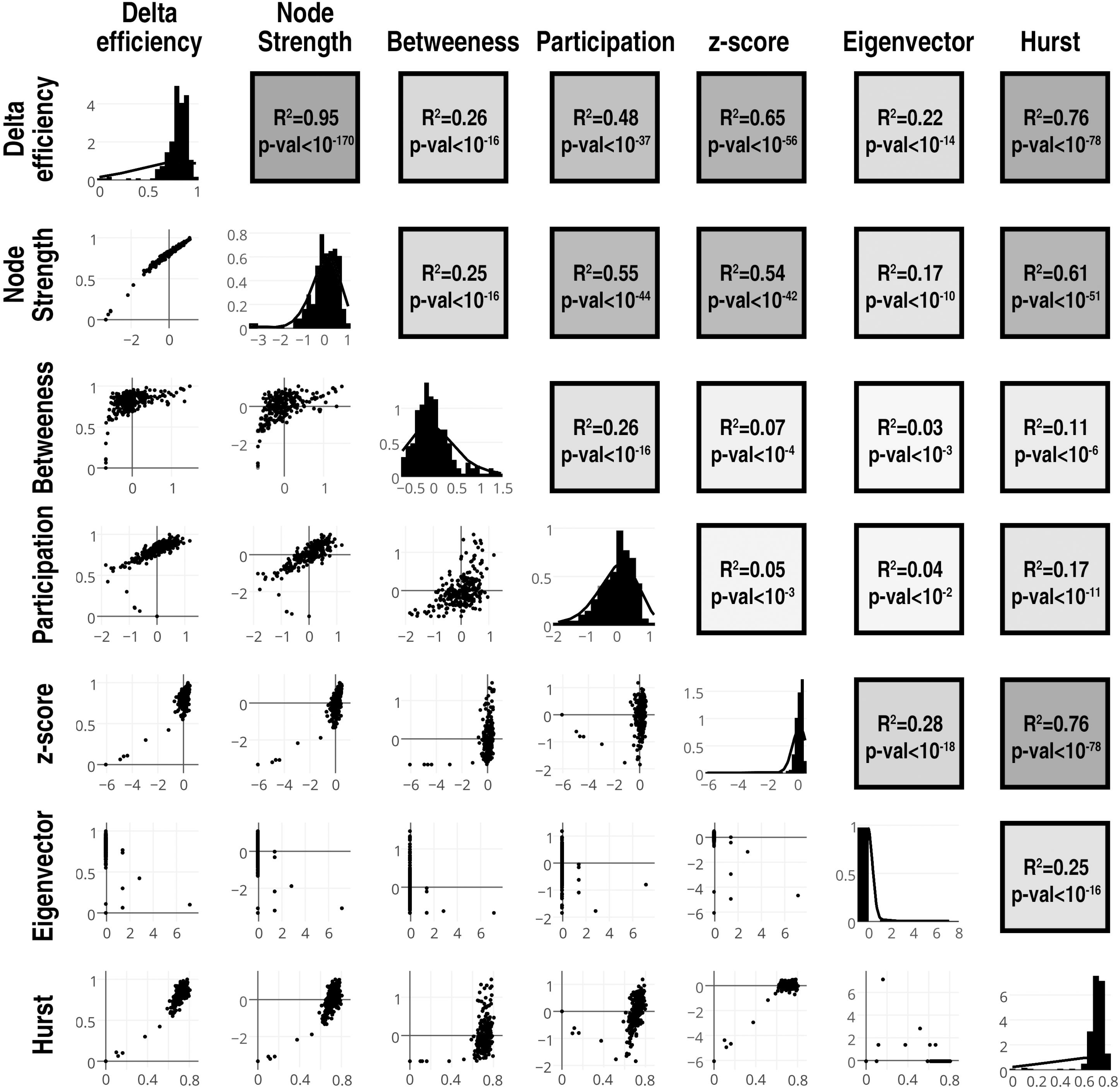
network measures compared to Hurst exponents. Plotmatrix of selected network centrality measures and H. Diagonal elements represent the histograms of each value after z-transforms. Above diagonal elements represent the corresponding R^2^ and p values as a heatmap.

